# Measuring social networks in primates: wearable sensors vs. direct observations

**DOI:** 10.1101/2020.01.17.910695

**Authors:** Valeria Gelardi, Jeanne Godard, Dany Paleressompoulle, Nicolas Claidière, Alain Barrat

## Abstract

Network analysis represents a valuable and flexible framework to understand the structure of individual interactions at the population level in animal societies. The versatility of network representations is moreover suited to different types of datasets describing these interactions. However, depending on the data collection method, different pictures of the social bonds between individuals could a priori emerge. Understanding how the data collection method influences the description of the social structure of a group is thus essential to assess the reliability of social studies based on different types of data. This is however rarely feasible, especially for animal groups, where data collection is often challenging. Here, we address this issue by comparing datasets of interactions between primates collected through two different methods: behavioral observations and wearable proximity sensors. We show that, although many directly observed interactions are not detected by the sensors, the global pictures obtained when aggregating the data to build interaction networks turn out to be remarkably similar. Sensors data yield moreover a reliable social network already over short timescales and can be used for long term campaigns, showing their important potential for detailed studies of the evolution of animal social groups.

## Introduction

Interactions between individuals are the foundation of complex social structures in human and other animal societies. Network analysis represents a valuable framework to understand the structure and evolution of these interactions, as it encodes a whole hierarchy of patterns, from individual-level interactions to complex population-level social structures. [1–3, 3–8].

With the increasing deployment of digital devices, new ways of collecting data, combined with new network analysis tools, have made possible the development of quantitative measures of these relationships and patterns in modern human societies, leading to the emergence of computational social science in the last decades [9]. For instance, social relationships have been inferred and studied using various data sources ranging from phone calls [10], e-mails [11, 12], online interactions [13], to face-to-face interactions measured by wearable sensors [14–20].

The availability of large volumes of data with high temporal resolution has thus contributed to the rapid expansion of data-driven computational studies of human relationships and human social networks. On the contrary, the data collection remains more challenging in the field of animal studies, because the data on animal interactions are still largely obtained from direct observations [6, 21]. Data resulting from such observations are extremely valuable as they often include detailed information about the nature, duration and location of the interactions between individuals. They thus allow researchers to grasp and investigate complex social patterns in animal groups. Unfortunately, observations are time costly and, if they are not performed for a long enough time per individual, suffer from a strong sampling effect [22]. Moreover, observations are almost always biased to some extent because, for example, the visibility of animals is not uniform and some interactions are more easily defined and recognized than others (see e.g. [22]).

Recently, a number of technological developments have started to be adapted and implemented to gather high-resolution behavioral data on non-human animals, leading to the adoption of the term reality mining, widely used in computational social sciences for the study of human social behavior and relations [23], to the case of non-human animal societies [24]. Machine-sensed data concerning the behaviour of animals can indeed now be collected and, most importantly, analysed. We refer to [25] for an overview of existing and emerging technologies used to collect data on movements, behaviour and interactions within animal groups. In particular, image based tracking software and machine learning tools can be used to identify and track animals and their trajectories from video data [26–29]. High-resolution GPS can also be used to analyze animals’ relative movements: for instance, GPS tracking of wild baboons revealed that a process of shared decision-making governs baboon movements [30]. Different types of data can also be collected jointly (as in the Sociometers deployed in human groups [31]), such as for instance GPS and audio recordings to investigate the role of vocalisation on the cohesion of a group of animals [32].

These recent developments also include proximity logging technologies based on wearable sensors that are able to provide information either on the distance between the sensor and a fixed receiver [33], or on the distance between two sensors [34–39]. These efforts have enabled the collection of high-resolution data sets in various contexts. Using these techniques presents a number of advantages. First, wearable sensors afford an objective and reliable definition of contact as a proximity event. Second, all individuals equipped with a sensor are monitored together, continuously and potentially for a long time without the need for constant human supervision. This enables in principle the collection of large data sets covering long periods of times and, consequently, makes it possible to investigate the evolution and stability of social relationships and social groups on long timescales. On the other hand, wearable sensors do not yield information on the type of behavioral interactions and they do not register contacts with individuals not wearing any sensor, such as very young individuals or out-group members for instance. The quality of the collected data might also depend on infrastructure constraints and potential technical failures, so that sampling issues must also be considered carefully [20, 40].

Thus, data obtained from direct observations and from wearable sensors infrastructures have very different nature and could in principle lead to very different descriptions and understanding of the social bonds between individuals and of the resulting social networks [41, 42].

Understanding to what extent the way of collecting data influences the final image of the social network, which elements affect this result and how, is therefore essential to assess the reliability of the outcomes of social studies. Due to the difficulty in collecting data using different methods at the same time and in the same population, few studies have been able to address these issues. In human groups for instance, comparisons between contacts registered by wearable sensors and in diaries have shown both similarities and differences between the data collected by these two methods. In particular, many contacts registered by sensors are not reported in surveys, especially for short contacts, while long contacts are better reported [43–45]. Comparison between sensors and direct observations or videos have yielded mixed results [46, 47]. Among animals, different types of networks built from the same data set of direct observations have been shown to differ [41, 42], while a social network deduced from co-presence in cognitive testing booths has been shown to correlate with the one obtained from directly observed interactions [48, 49]. However, we are not aware of studies using data collected in the same population with on the one hand wearable sensors and on the other hand direct observations.

Here, we address this issue by collecting, describing, analyzing and comparing two data sets based on dyadic interactions between individuals belonging to a group of Guinea Baboons (*Papio papio*). The data span a time-window of almost one month between June and July 2019 and were collected through two different methods: (i) behavioral observations by trained human observers and (ii) an infrastructure based on wearable sensors (see http://www.sociopatterns.org/).

For these two data sets, we first test the agreement between observations and the sensors data at the level of single events: we systematically check whether an observed interaction was also registered by the sensors. Overall, only a limited fraction of observed interaction events were registered, with strong fluctuations depending on the day of observation and type of behavior. However, and despite this poor agreement at the level of single events, we show that the time-aggregated networks, and hence the pictures of the group social structure, are remarkably similar.

Finally, we analyse the amount of time that it takes using each data collection method to obtain a robust social network, by comparing the social networks obtained using different time aggregation windows. Strikingly, the social network obtained with just one day of sensor data is very similar to the one based on the aggregation over one whole month of data. Comparatively, the network obtained from observations fluctuates more between short and long aggregation windows because of stronger sampling effects. This shows the potential of wearable sensors infrastructures to detect changes in a social group organization on short timescales and to also monitor its long term evolution.

## Methods

### System setting and data collection

The data collection involved a group of captive Guinea baboons (*Papio papio*) living in an enclosure of the CNRS Primate Center in Rousset-sur-Arc (France). The entire group consisted of 19 individuals (7 males and 12 females) with age ranging from 1 to 23 years old.

#### Behavioral observations

The behavioral observations were recorded between June 13^*th*^ and July 10^*th*^, 2019 using the focal-sampling method [50]. Observations were carried out for five days a week (from Monday to Friday) for a total of 20 days, with two sessions of ∼ 2 hours a day at different hours each day, ranging from 8am to 5pm. During each session, a trained observer focused on each individual for a period of 5 minutes and recorded its behaviors. The order in which the different individuals were observed was reshuffled at each session. Thirteen behavioral categories corresponding to interactions were recorded, namely: ‘Grooming’, ‘Presenting’ (Greeting),’Playing with’, ‘Grunting-Lipsmacking’, ‘Supplanting’,’Threatening’, ‘Submission’, ‘Touching’, ‘Avoiding’, ‘Attacking’, ‘Carrying’, ‘Embracing’, ‘Mounting’, ‘Copulating’, ‘Chasing’ (see Supplementary Material for details). Another behavior of interest was ‘Resting’. Resting is often considered as a behavior performed in isolation, i.e., does not correspond to an interaction between individuals. In some cases however, when two or more individuals were resting together at less than one meter from each other, this was considered as ‘social resting’ and also counted as an interaction. ‘Resting’, ‘Grooming’ and ‘Playing with’ include a duration and are called *State* events. The other types of behavior do not have a duration assigned and are called *Point* events. For each observed behavior, the individuals involved were recorded as well as the starting and ending time of each state event, and, for each point event, the time at which it took place.

In addition, two categories were included, namely ‘Invisible’ and ‘Other’, to refer to the cases in which the individual was not seen by the observer or the behavior was not included among those listed above.

#### Wearable sensors data

A subgroup of 13 baboons, consisting only of juveniles and adults (all individuals at least 6 years old) were collared with leather collars. The collars were fitted with wearable proximity sensors (RFID tags) developed by the SocioPatterns collaboration (http://www.sociopatterns.org/), already used in many studies involving humans [14, 16, 18, 20, 40, 44, 51], and recently also animals [39]. In our setting, each sensor was secured in a customized box specially designed with a 3-D printer to contain the sensor and a long-life battery connected to it. The boxes were positioned on the front side of the individuals (Fig 1).

**Fig 1.**
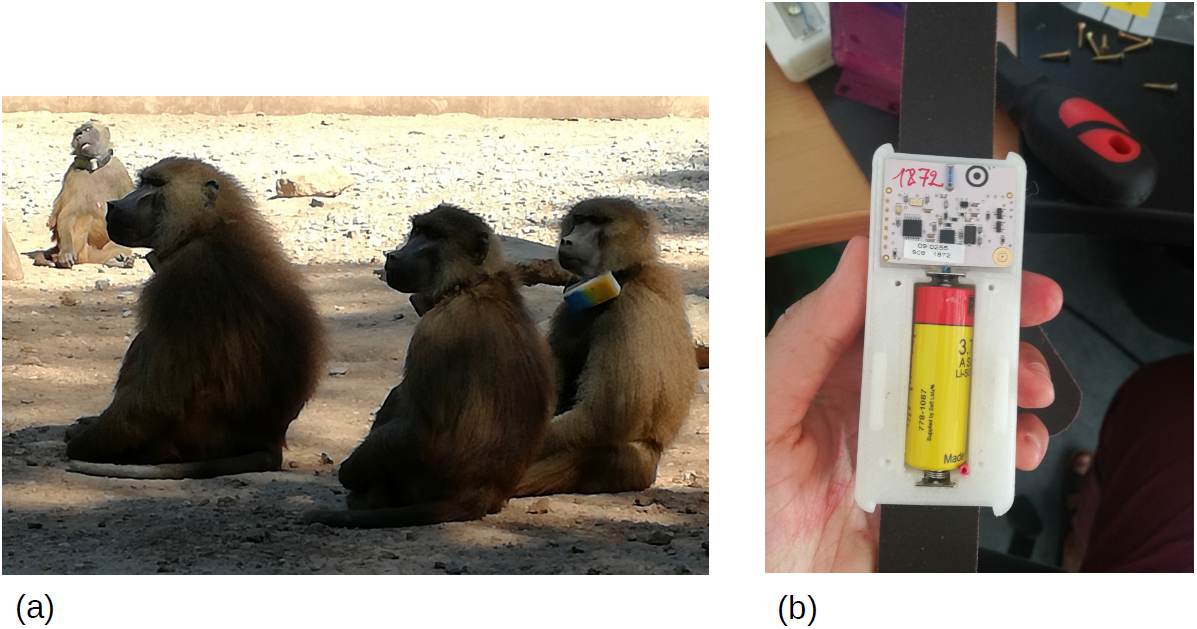
(a): Individuals in the enclosure, wearing collars with the attached boxes embedding the tags. (b): Interior of a single box containing the tag (top) and the connected battery (bottom)

The sensors exchanged low-power radio packets in a peer-to-peer fashion. Thanks to the very low power used, the reception by the sensor of an individual A of a radio packet emitted by the sensor of another individual B was a good proxy for a close proximity (≲1.5 m) of individuals A and B [14]. Moreover, the radio frequency emitted by the RFID tags was absorbed by body water, so that the radio packets tended to propagate mostly towards the front of the individual wearing the device. The packets exchange rate depended thus on the mutual orientation of the individuals and the infrastructure detected mainly face-to-face interactions. The detected spatial proximity relations were relayed from RFID tags to radio receivers (RFID readers), which were installed around the enclosure and connected to a local area network (LAN). A central server received the data, timestamping and storing each event.

Data were finally aggregated with a temporal resolution of 20 seconds (for more details see [14]): we thus defined two individuals to be in contact during a 20s time window if their sensors exchanged at least one packet during that interval, and the contact event was considered over when the sensors did not exchange packets over a 20s interval.

In the following, we will refer to the observed behaviors corresponding to interactions as ‘interactions’ or ‘observed interactions’, and to the contacts collected by the sensors as ‘contacts’ or ‘contact events’.

For the first ten days of data collection (June 13^*th*^ -23^*rd*^) only two readers were installed around the enclosure, whereas in the successive days a third reader was added to ensure a better coverage. The data collection went on even after the observation period was over and is on-going at the time of writing of this paper. We consider here mainly the data collected between June 13^*th*^ and July 10^*th*^, 2019, i.e., during the period of the observations, and use also the data collected afterwards and until August 27^*th*^ to assess stability over longer time scales.

### Ethics Statement

The baboons lived in an outdoor enclosure (700*m*^2^) connected to an indoor area that provided shelter when necessary. Water was provided ad libitum within the enclosure, and they received their normal ratio of food (fruits, vegetables, and monkey chow) every day at 5 pm. The baboons were all born within the primate centre. This research was carried out in accordance with French standards and received approval from the national French ethics committee, the “Comité d’Ethique CE-14 pour l’Expérimentation Animale” (approval number APAFIS#4816-2015091110584769).

### Data analysis

#### Comparison of the two data sets at the level of single events

We compute the fraction of observed interactions that were also detected by the wearable sensors infrastructure system as follows. Each observed interaction event involves two individuals *i* and *j* and is assigned a time *t* (for point events) or an interval [*t*_*start*_, *t*_*stop*_] (for state events), with *t*_*start*_ and *t*_*stop*_ the moments of beginning and end of the interaction, respectively. Note that the observed interactions are often directed, with an actor individual and a recipient individual. However, since the proximity events registered by the wearable sensors are not directed, we consider undirected versions of the behavioral data (i.e., the direction of the interaction is not taken into account).

An observed interaction event is then considered as *tracked* if at least one RFID packet was exchanged between the sensors of individuals *i* and *j* within the time window [*t* − Δ*t, t* + Δ*t*] or [*t*_*start*_ − Δ*t, t*_*end*_ + Δ*t*], for point and state events respectively, where Δ*t* is a tolerance interval. This tolerance is introduced to take into account three elements: (i) the potential delay of the observer in reporting the interaction with respect to its actual occurrence; (ii) the 20s time aggregation of the RFID sensors data (iii) the possible asynchrony between the observer’s tablet computer in which observed interactions were registered and the time of the computer storing the sensors’ data.

Obviously, to compute the fraction of tracked interaction events, we only consider observed events involving two individuals wearing collars embedded with sensors.

#### Comparison of the resulting aggregated networks

For each data set, we can construct on any temporal window an aggregated network in which nodes represent individuals and weighted links give a summary of the recorded contacts or observed interactions during that time window. We first consider as time window the whole period in which observations were carried out, and we restrict the observational data to the 13 individuals with collars and sensors. We thus obtain a “contact network” from the wearable sensor data and an “interaction network” from the observed interactions, both covering the period from June 27^*th*^ to July 10^*th*^, 2019. Both networks are undirected and weighted.

In the aggregated interaction network, a weighted link between nodes *i* and *j* is drawn if at least one interaction was observed between *i* and *j* during the aggregation time window. The weight 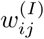 of the link between individuals *i* and *j* is given by the total number of interaction events recorded between *i* and *j* during this time. Note that we use here the number of interactions and not their total duration in order to account also for point events. Similarly, in the contact network, a link was drawn between *i* and *j* if at least one contact was recorded between them by the sensors infrastructure, and the corresponding weight 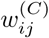 is given by the number of contacts recorded by the sensors between *i* and *j*.

We compare the contact and interaction networks using several metrics. We computed the Pearson and Kendall’s *τ* correlation coefficients between the two lists of weights to measure respectively the linear correlation and the similarity of the orderings of the weights in the two networks. We also considered two different versions of cosine similarity measures between the networks weights. A cosine similarity measure is in general defined between two vectors, and is bounded between −1 and +1. It takes the value −1 if the vectors are proportional with a positive proportionality constant, a value of 1 if the proportionality constant is negative, and 0 if they are perpendicular. For positive weights as in our case, it is bounded between 0 and 1. We consider first a Global Cosine Similarity (GCS) measure between the two networks as the cosine similarity between the two vectors formed by the list of all links weights in each network (using a weight 0 if a link is not present):

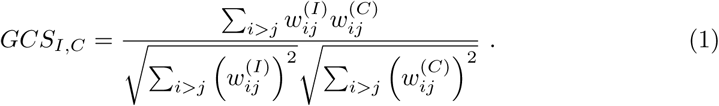

We moreover consider local versions of the cosine similarity: the Local Cosine Similarity (LCS) of a node *i* is given by the cosine similarity between the vectors of weights involving *i* in each network:

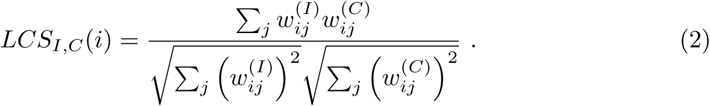

*LCS*_*I,C*_ (*i*) is thus equal to 1 if *i* has been detected as linked with the same individuals in the two data sets with proportional weights.

It is equal to 0 if *i* has disjoints sets of neighbours in the two networks. Here we use the average LCS value over all individuals as a measure of similarity between the two networks.

To get a better grasp of the values obtained, we consider a null model in which the weights are reshuffled among the links for one of the networks. We perform 1000 realizations of this reshuffling and recompute the values of correlations and similarities in each realization, obtaining a null distribution for each measure.

#### Other aggregation timescales: fluctuations and convergence

For each data set, the aggregation procedure yielding an aggregated network can be performed on any time window. In addition to the whole observation time windows, we consider shorter time windows of 1 day, 3 days and one week. We then study the stability and dynamics of each network by computing for each data set the cosine similarities between the networks aggregated on all pairs of time window (with a given length). For instance, if we consider two different time periods *t*_1_ and *t*_2_, we denote by 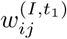 and 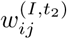 the weights of the links between individual *i* and *j* in the interaction networks aggregated over *t*_1_ and *t*_2_ respectively, and the local cosine similarity of *i* in the interaction network between *t*_1_ and *t*_2_ is

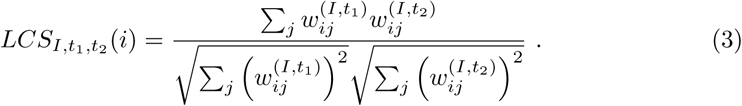

Local and global cosine similarities can be defined in the same way for each data set, and can also be defined between networks aggregated on time windows of different lengths. We compute for each data set similarities between the network aggregated on the whole observation time window and networks aggregated on the first *n* days of observations, in order to understand how fast the weight structure of each network converges to its fully aggregated version.

## Results

### Single interaction and contact events

#### Behavioral data

The total number of behaviors recorded for the entire group of 19 individuals is 5, 377. From this full data set of observations we keep just the behaviors involving the 13 individuals that were carrying collars with wearable sensors. Among the 995 observed interactions regarding this juvenile/adult subgroup, 944 (∼ 95%) are affiliative social behaviors (Grooming, Resting, Presenting, Grunting-Lipsmacking, Touching, Mounting, Embracing, Playing with) which are the most relevant to this study. Moreover, grooming and (social) resting represent more than 98% of the state events and ∼ 65% of the total.

Figure 2 represents the distributions of durations for the observed interactions (i.e., with associated duration) both for the subgroup of collared individuals and for the whole group, which includes very young juveniles and babies. Durations cover a broad range of values with a cut-off point at 300s (5 minutes) corresponding to the duration of the focal observation (i.e., some interactions lasted more than 300s but their total duration is unknown). We also note that the distributions of events concerning the whole group or only the collared individuals have similar shapes, with however more short interactions when babies are included.

**Fig 2.**
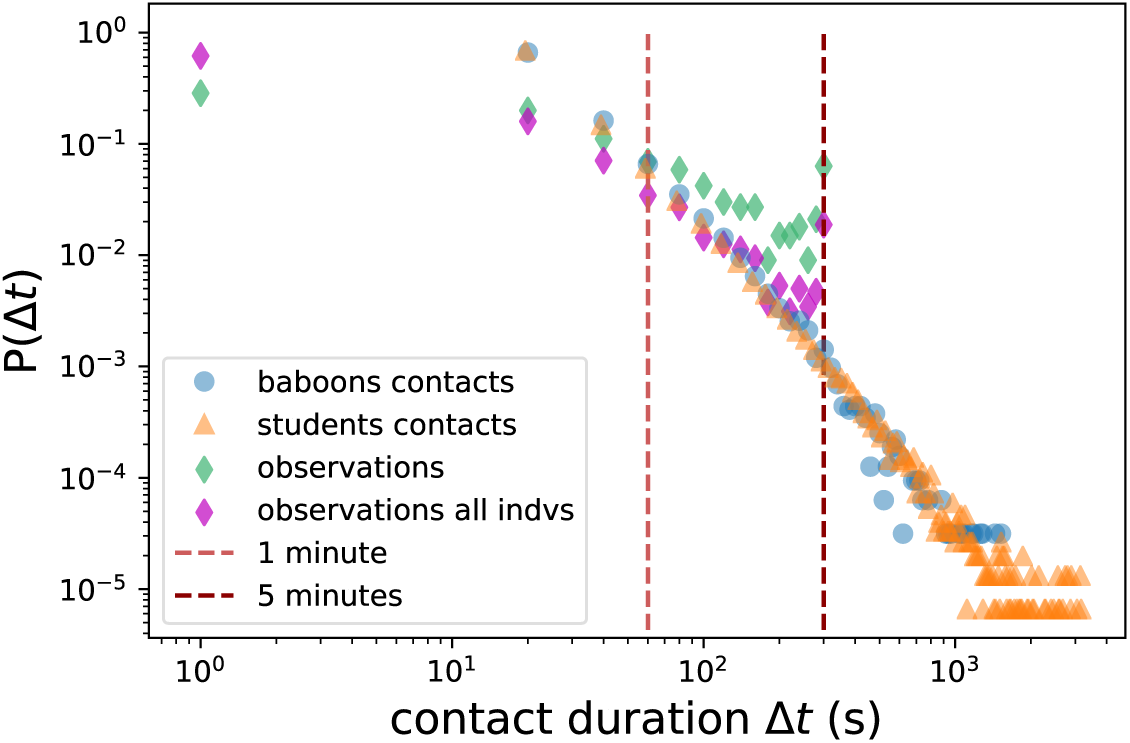
Durations of contacts and observed interactions. Distributions of durations for: contacts detected by the wearable sensors infrastructure (blue dots; average = 39.6s, std = 52.7s, median = 20.0s, max = 1, 520s); contacts between students in a school in Utah (USA) measured by an infrastructure based on wireless ranging enabled nodes (WRENs) [19] (orange triangles; average = 39.6s, std = 72.8s, median = 19.5s, max = 3164s); state events in observed interactions (green diamonds for interactions involving only the 13 collared individuals; average = 81.3s, std = 90.3s, median = 41.0 s, max = 300s; magenta diamonds for all interactions; average = 43.8s, std = 64.6s, median = 18.0 s, max = 300s). Note that according to the observational method used in this study, individuals are observed for 5 minutes (300 seconds) at a time. The peak value at 300s for the observations data is therefore an artifact of the observation method.

#### Contacts registered by the sensors

During the period in which observations were carried out, 31, 783 contact events were recorded between the 13 individuals. Of these, 4, 823 (15% of the total) were recorded during the periods of behavioral observations. The number of contacts per day was ∼ 1, 135 on average and ranged from 754 (June 28^*th*^) to 1, 768 (June 13^*th*^) for a total duration of 1, 259, 500 s (349 hours). Contact durations varied on a very broad range: most contacts were short, with an average duration of 39.6s, and 95% of the contacts lasted less than 2 minutes, but contacts as long as 1, 520s (∼ 25 minutes) were recorded, and the contact durations form a continuous distribution spanning all values in between (see Fig. 2), as observed in many different contexts for human and animal groups [14, 15, 18, 20, 39, 44]. In fact, we report on the same graph the statistics of contact durations measured by wearable sensors between students in a school, reported in [19] and freely accessible: it turns out that the distributions of contact durations of baboons and of humans are indeed very similar.

#### Comparing interactions and contacts

Figure 2 allows to compare the distributions of durations of the observed interactions and of the contacts registered by the sensors infrastructure. Although they differ, with in particular a distribution extending to much larger values for the contacts, they are both broad and spanning a large range of values. The limited range of the interactions durations is due to the observational protocol, since all durations above 300 seconds are cut-off at that value. In addition, the duration of interactions starting before the start of a given observation 5-minutes time-window, or ending after its end, are necessarily underestimated, which biases the resulting distribution in a complex way. Overall, it seems possible that the two distributions would have similar slopes at large durations if this cut-off were not enforced although a detailed study of the biases introduced by the cut-off is beyond the scope of this work.

To go beyond this statistical comparison, we perform a detailed matching procedure, as described in Methods, between each single observed interaction (in the behavioral data) and the contact events (obtained from the sensors data). Table 1 gives the results of this matching, for different categories of interactions and different values of the tolerance Δ*t*. The fraction of observed interactions finding a match in the sensors data is quite low, with a slightly better performance when the tolerance is increased. For Δ*t* = 20*s*, on average only one third of the observed interactions appear in the data obtained from the sensors infrastructure. The fraction is notably larger if we consider only grooming events, which are known to be very important socially [52]. We note that in this case the fraction of tracked observed interactions is about 50%, a value very close to the one recently obtained in [47] in a comparison between the contacts between human individuals as observed in an annotated video and as registered by wearable sensors. On the contrary, the fraction is lower for very short events such as greetings.

**Table 1.** Fractions of observed interactions with a corresponding match in the sensors data. We consider an interaction to have a match in the contacts data if the pair of individuals involved in the interaction appears in the sensors data in the same time window ±Δ*t* (see Methods). We report the overall fraction of matched interactions (first row), the average fraction over the days (second row) with the corresponding standard deviation (third row), and the minimum and the maximum (fourth and fifth row) fractions of tracked interactions over the different days. The values were computed for different delay parameters Δ*t*, and considering either all interactions, only state events (i.e., interactions with duration), only grooming events and only greeting events.

|  | All Interactions |  |  | State Events |  |  | Grooming |  |  | Presenting (Greeting) |  |  |
| --- | --- | --- | --- | --- | --- | --- | --- | --- | --- | --- | --- | --- |
| $\Delta t$ | 20s | 40s | 60s | 20s | 40s | 60s | 20s | 40s | 60s | 20s | 40s | 60s |
| Tot. | 0.32 | 0.36 | 0.39 | 0.36 | 0.4 | 0.43 | 0.48 | 0.53 | 0.56 | 0.23 | 0.27 | 0.29 |
| Avg. | 0.32 | 0.36 | 0.38 | 0.37 | 0.4 | 0.42 | 0.49 | 0.53 | 0.55 | 0.2 | 0.24 | 0.27 |
| Std. | 0.12 | 0.13 | 0.13 | 0.12 | 0.13 | 0.14 | 0.19 | 0.21 | 0.22 | 0.2 | 0.2 | 0.21 |
| Min. | 0.1 | 0.1 | 0.12 | 0.11 | 0.11 | 0.14 | 0.17 | 0.17 | 0.17 | 0.0 | 0.0 | 0.0 |
| Max. | 0.54 | 0.63 | 0.63 | 0.61 | 0.64 | 0.64 | 0.83 | 0.83 | 0.89 | 0.67 | 0.67 | 0.83 |

We finally note that we can consider the reverse procedure, considering the contacts registered by the sensors as ground truth. To this aim, we restrict the sensors’ contact data to the time windows corresponding to the behavioral observation periods: only 6.63% of these contacts were recorded by the observer as interactions. Note that this small number is not surprising, as the observer focuses on one individual at a time, while the sensors’ infrastructure registers contacts between all collared individuals during the same time window.

### Comparing interaction and contact networks

Both interaction and contact networks, obtained from the aggregation over the whole period of observation, are very dense, with respectively 70 and 78 links (in particular, the contact network is fully connected, i.e., with at least one contact registered between all pairs of individuals).

The two networks have by nature widely different weights, due to the differences in the methods of measurement. In particular, the number of observed interactions is strongly limited by the amount of time dedicated to the observation of each individual. As a result, the number of observed interactions between a given pair of individuals is at most of a few tens. On the other hand, sensors are active at all times and the weights of the contact network span several orders of magnitude, as common in such data sets [14]. Despite these differences in the range of weights of the two networks, Figure 3 show that the statistical distributions have in fact very similar shapes.

**Fig 3.**
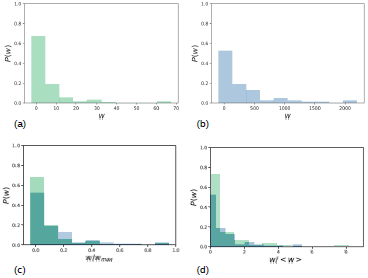
Distributions of weights. Probability distributions of interaction (a) and contact (b) networks weights. Despite the very different values of the weights, their distributions have similar shapes. In panels (c) and (d) the same distributions are shown after rescaling the weights of each network by either the maximum weight (c) or the average weight (d).

In addition, Figure 4 gives a visualization of the two weighted networks in which the weights have been rescaled to have comparable widths. This figure highlights some important similarities in the structure of the strong links of both networks: relevant examples include the links Kali-Pipo (female-male) and Angele-Felipe (female-male), as well as the triad Atmosphere-Harlem-Violette (female-male-female) with two similarly strong links Atmosphere-Harlem and Harlem-Violette, and a very weak link Atmosphere-Violette.

**Fig 4.**
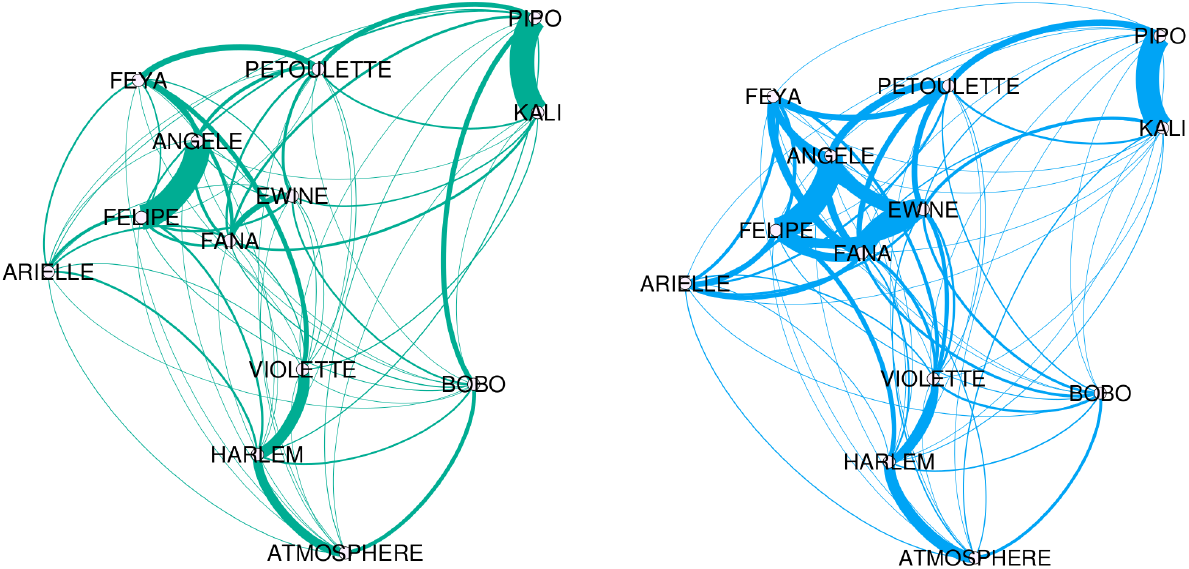
Graphs visualization. Visualization realized with Gephi software (www.gephi.org, see also [53]) of the interaction network (left) and the contact network (right), aggregated over the entire observation period. The thickness of the lines is proportional to the links weights (scaled in order to have comparable links widths in both networks). The figure shows in a qualitative way the high resemblance between the patterns of strong and weak links of the two networks.

To go further, we present in Table 2 a systematic comparison of the strongest links in both networks. The table reports the lists of the ten strongest links in each network. The two strongest links are the same in both networks, and more than half (6/10) of these links appear in both interaction and contacts networks. The links that are among the top ten of one network but not of the other are moreover within the top 20 strongest links of the other network, except for one. The exception is the only link between two adult males appearing in Table 2, namely the link BOBO-PIPO, ranked 8^*th*^ for the interaction network but only 55^*th*^ in the contact network. The interactions between adult males are usually short greetings (‘presenting’), which are not face-to-face but face to rear (see the description of behaviors in the Supplementary Material), making this interaction harder to be detected by the sensors. We checked that this was indeed the case for the BOBO-PIPO link, with almost only greetings and other point events (92%).

**Table 2.** Top 10 strongest links. For each network, links are ordered based on their weights, from the strongest to the weakest. The links in bold are the ones present in both top ten rankings.

| Rank | Interaction Network (observations) | Contacts Network (sensors data) |
| --- | --- | --- |
| 1 | **FELIPE - ANGELE** | **FELIPE - ANGELE** |
| 2 | **KALI - PIPO** | **KALI - PIPO** |
| 3 | **VIOLETTE - HARLEM** | **FANA - EWINE** |
| 4 | **ATMOSPHERE - HARLEM** | EWINE - ANGELE |
| 5 | FEYA - PETOULETTE | **ATMOSPHERE - HARLEM** |
| 6 | **FANA - EWINE** | EWINE - FELIPE |
| 7 | PETOULETTE - PIPO | **VIOLETTE - HARLEM** |
| 8 | BOBO - PIPO | **FANA - FELIPE** |
| 9 | **FANA - FELIPE** | FANA - ANGELE |
| 10 | EWINE - PETOULETTE | FEYA - ANGELE |

Finally, as described in Methods, we use four different indicators to give a quantitative estimation of the similarity between the networks. Each indicator is computed for the two empirical networks and for 1000 realizations of the null model described in Methods, and we compare in Figure 5 the empirical value and the distribution of values obtained with the null model. The global and average local cosine similarity are extremely high (close to 0.9), as well as the Pearson correlation coefficient (0.83), while the Kendall rank correlation coefficient is still large (0.58) but more impacted by the large number of links with low weights, whose order is expected to be less stable than the one of the strong links. In all cases, the empirical values lie far above any value obtained in the null model realizations, as clearly seen on the figure (see also the Supplementary Material for a scatterplot of the weights of the links in the two networks and for the values of the local cosine similarities of each individual).

**Fig 5.**
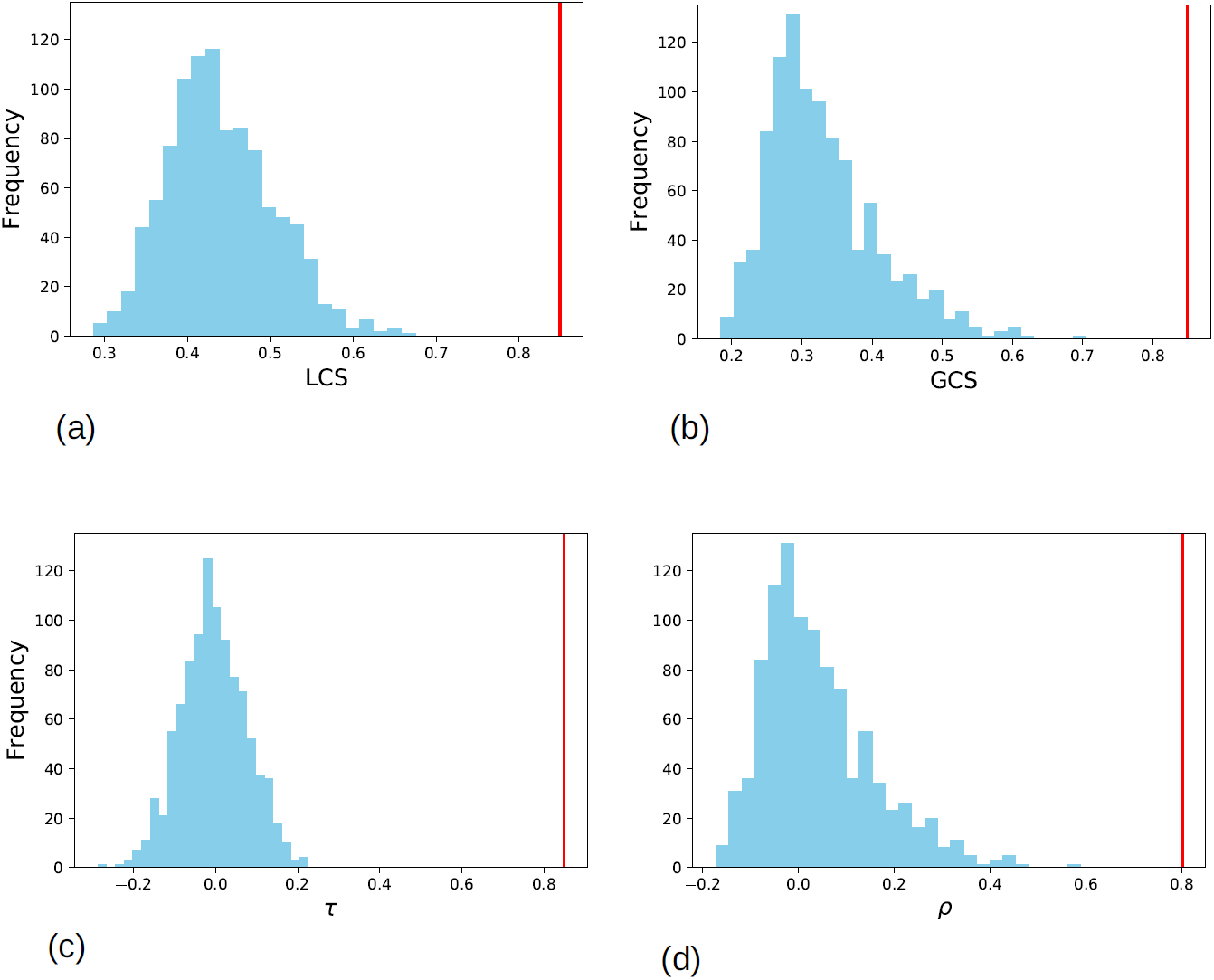
Similarity metrics. Quantitative comparison between the contact network and the interaction network through several correlation and similarity measures. In each panel the empirical value is presented as a vertical red line, together with the distribution of 1000 values (light blue) obtained using a null model in which the weights of the contacts network were reshuffled and reassigned at random to the links. (a) average of the local cosine similarity, i.e. average of the cosine similarity between the ego-networks of single nodes (empirical value: 0.91, distribution: average = 0.44, std = 0.06); (b) global cosine similarity between the lists of weights of the two networks (empirical value: 0.84, distribution: average = 0.33, std = 0.08); (c) Kendall rank correlation coefficient between the weights of the two networks (empirical value: 0.58, distribution: average = −0.004, std = 0.080); (d) Pearson correlation coefficient between the weights (empirical value: 0.80, distribution: average = 0.04, std = 0.11).

### Other aggregation timescales

As described in Methods, we also considered other timescales on which to aggregate the data coming from the observations and from the sensors infrastructure. Indeed, the time constraint of observational measures implies that the amount of information concerning each individual per day is relatively small. Building a reliable image of the social bonds between individuals requires thus many days of observation and an aggregation window of one month is usually advocated [22, 48]. In the case of data collected through wearable sensors on the other hand, a large amount of contacts is already recorded after a few hours. However, it is a priori unclear whether the structures present on short timescales such as e.g. one day of data fluctuate from day to day or are stable and already representative of the group social structure and of its strong links. Indeed, the structures present in the monthly aggregated network might result either from a superposition of different daily networks, or, on the contrary, from the repetition of the same contact network every day.

To investigate this issue, we compute aggregated interaction and contact networks over different timescales (1 day, 3 days, 1 week), obtaining in each case series of successive snapshots corresponding to the interactions observed or to the contacts measured in successive time windows. For each type of network, we compute the cosine similarities (local and global) between each couple of snapshots to determine how stable networks are, once aggregated on such timescales. We show in Fig. 6 the resulting color-coded matrices for the average local cosine similarity: the values obtained for the interaction network are much lower than for the contact network, showing that the former fluctuates much more than the latter. We show in the Supplementary Material that the interaction networks aggregated at daily scale are even more fluctuating, and that important differences are measured even between weekly aggregated successive networks. On the other hand, daily networks obtained from sensor data are already quite stable.

**Fig 6.**
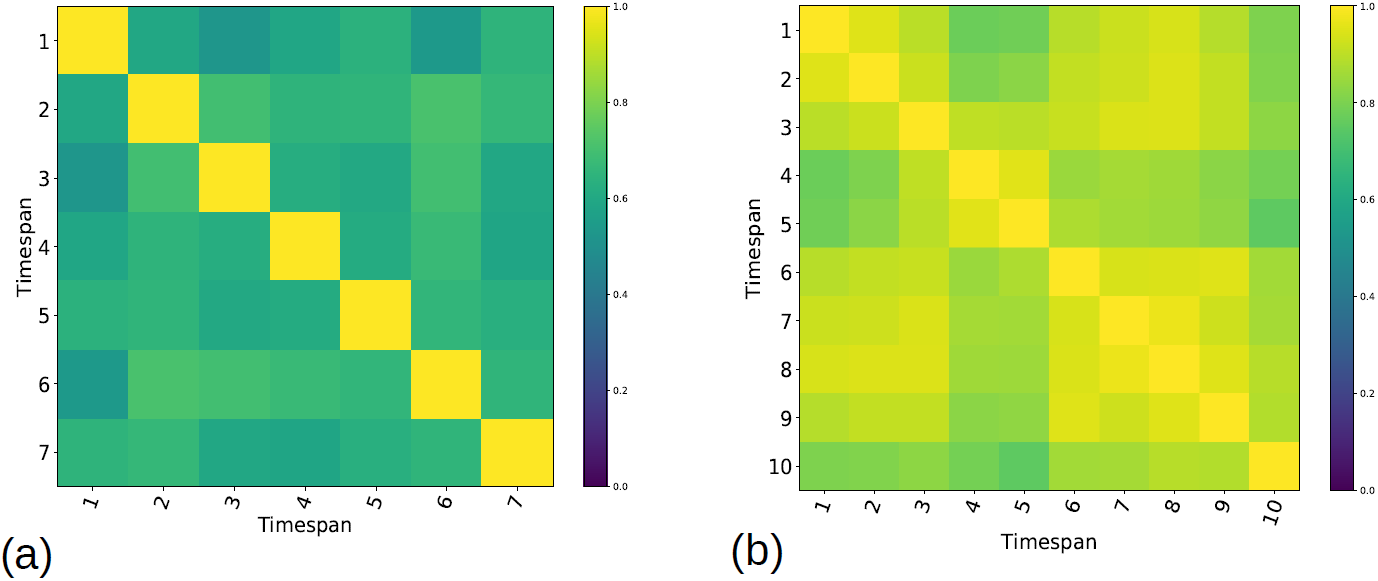
Cosine similarities between 3-days networks. Color-coded matrices of average local cosine similarity values between every couple of 3-days (a) interaction networks (min = 0.52; mean = 0.63) and (b) 3-days contact networks (min = 0.75; mean = 0.89). A smaller number of snapshots is obtained for the interaction network because no observations were carried out during the week-ends.

We then build interaction and contact networks on time windows of increasing lengths, starting from the beginning of the observations (using only the first day of data, then the first two days and so on), and compare them with the aggregated network based on the whole observation time window. Figure 7 shows the resulting GCS and average LCS as a function of the length of the time window considered. The obtained similarities are already close to 1 when only one day of the sensors data is used (GCS = 0.93, Avg. LCS = 0.90), and remain at high values for longer time windows. In fact, the inset shows that similarity values rise above 0.8 as soon as about 10 − 12 hours of data are collected. Comparatively, the similarities with the final aggregated network increase much more slowly for the interaction data than for the contact data. For instance, the one-day interaction network has much smaller similarity with the monthly one (GCS = 0.68, Avg. LCS = 0.58). However, they reach very high values already after 9 − 10 days of observation.

**Fig 7.**
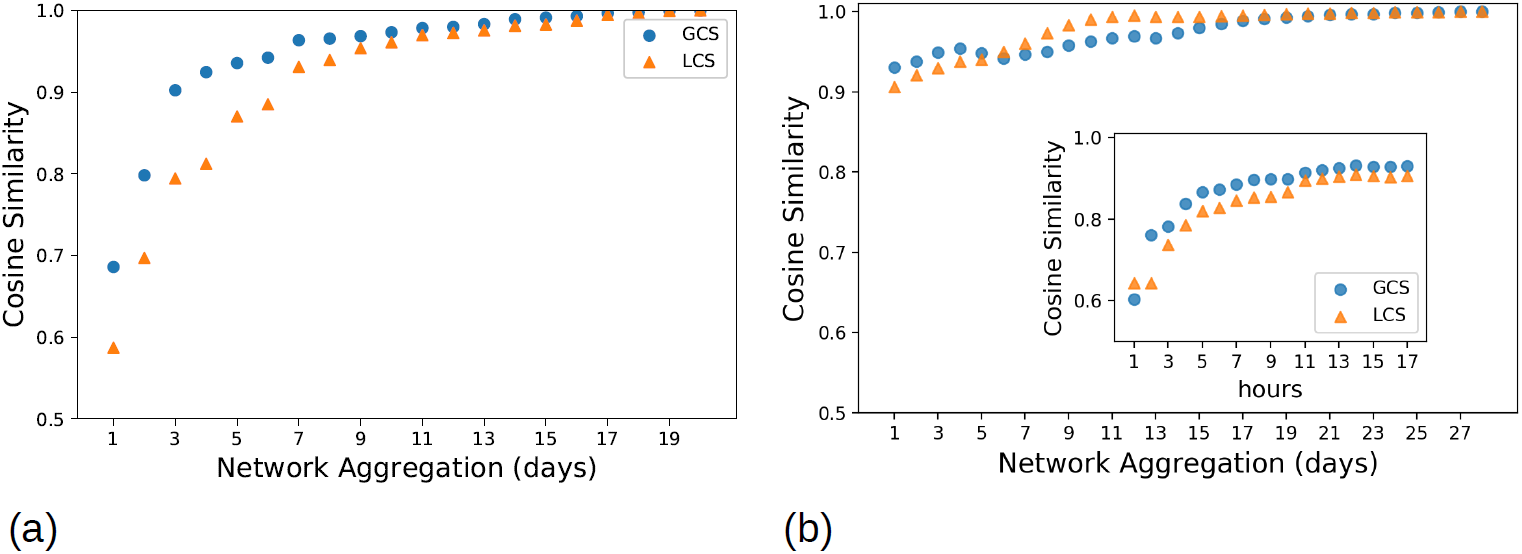
Global and average local similarities between the networks aggregated on the whole period of observation and networks aggregated on shorter time windows. (a): interaction networks; (b): contact networks.

Finally, to illustrate the possibility to explore long timescales using the sensors infrastructure, we consider the data collected after the observation period was over. Figure 8 shows the global and average local cosine similarities between networks aggregated on weekly timescales, from June 13^*th*^ to August 27^*th*^, 2019. The picture emerging from Fig 8 is the one of a very stable network, with average similarity values of 0.89 and 0.87 for the average LCS and for the GCS, respectively. Note that the minima of the similarities between weekly networks (0.75 and 0.61) are observed for the week July 18^*th*^-25^*th*^, in which the infrastructure actually failed for a couple of days, resulting in data loss for July 20^*th*^ and 21^*st*^.

**Fig 8.**
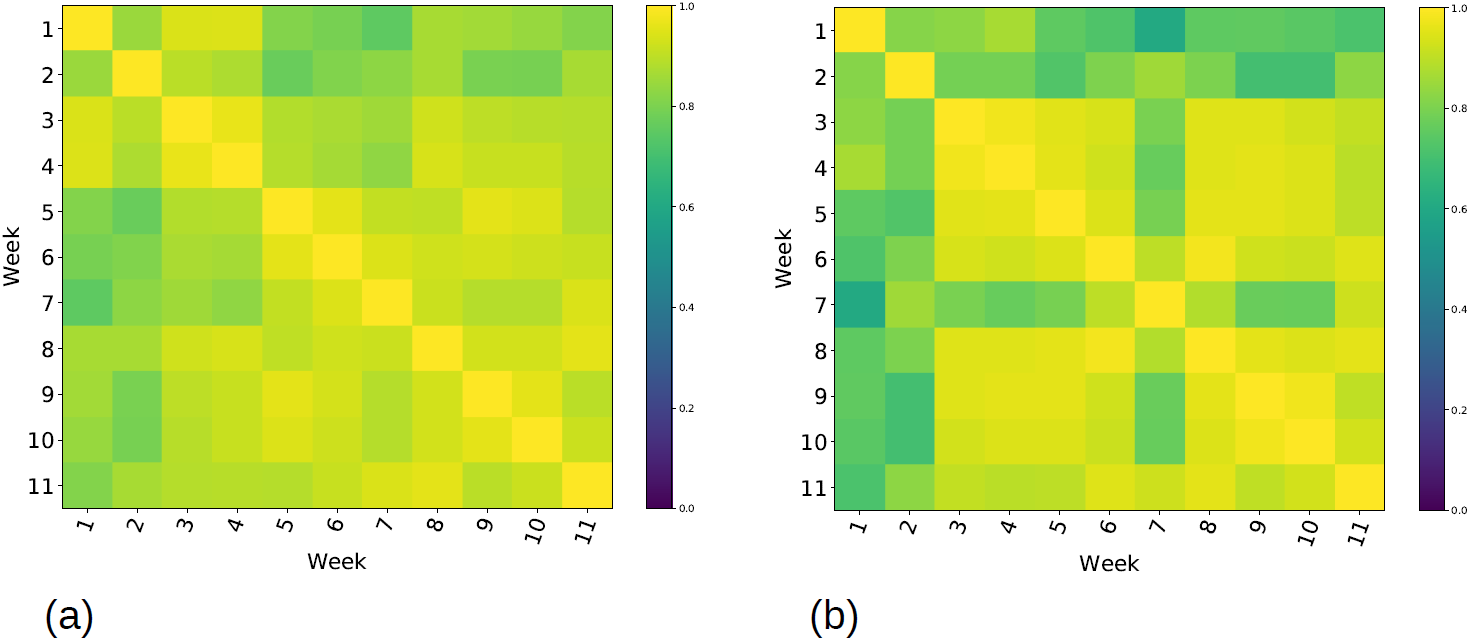
Stability of weekly-aggregated contact networks on long time scales. Color-coded matrices of the average local cosine similarity values (a) and global cosine similarity values (b) for all pairs of weekly contact networks from June 13^*th*^ to August 27^*th*^.

## Discussion

In this paper we analyzed and compared data sets describing social relationships in a group of non-human primates, collected through two different methods: behavioral observations and wearable proximity sensors. Sensors and behavioral observations methods have different advantages and limitations that influence their ability to detect contacts and interactions between individuals. On the one hand, observational methods provide high-quality data, through which we can distinguish different behaviors and describe in depth social relationships between individuals. However, the data suffers from several sampling issues: only a limited amount of time can be spent observing each individual, and data may be completely absent on certain days for logistical reasons (the week-ends in our case). Biases related to the observation technique can also occur and are difficult to estimate. The cutoff on the duration of each individual observation leads moreover to an underestimation of the duration of long interactions, which can be particularly important to determine the social structure of a group. Finally, the total duration of the observation period is usually limited to a few weeks and the group can not be monitored continuously on very long time scales, for clear practical reasons.

On the other hand, sensors provide a large amount of data in a continuous manner, with a high temporal resolution and potentially on very long timescales. However, the advantage of having an objective definition of a contact as an exchange of radio packets between sensors is at the same time a limitation since no information on the behavior of the individuals in contact is available. In particular, it is not possible to distinguish affiliative behaviors from agonistic ones, and it is therefore not possible to determine the dominance hierarchy using only wearable proximity sensors. Moreover, two types of sampling biases need to be mentioned. First, the directionality of the sensors limits the detection to approximately face-to-face interactions, while some social interactions between primates (in particular, greetings or social resting) occur when individuals have different mutual orientations. Second, individuals not wearing the sensors are by definition absent from the data. In the case of human populations, it has been shown that a uniform population sampling does not alter the statistical properties of the contact network between individuals [20]. However, the absence of data concerning a specific subgroup of individuals with behavioral patterns different from the rest of the group (such as all the individuals less than 6 years old, in our case) is a clear limitation. Another difficulty could also come from data losses whenever a part of the infrastructure fails. We show however in the Supplementary Material by simulating the failure of a reader that the structure of the social network deduced from the sensor data remains very stable even when the amount of data lost is important.

We have performed a comparison of the two data sets at various levels of detail. At the most detailed level, we have checked for each observed interaction whether a contact was registered by the sensors at the same time. This matching turns out to be very limited: about one third of the observed interactions was also recorded by the sensors, but this amount fluctuates and depends strongly on the type of interaction. In particular, for interactions that tend to last, such as grooming, the percentage of tracked events rises to ∼ 50% and is above ∼ 80% in some days of observations. This is particularly important as grooming behavior is for primates one of the core social interactions allowing to define the social structure of the group [52]. For short and elusive interactions instead, like greeting, this percentage is only of about ∼ 20%, which can be explained by the fact that greetings among primates are most often not face-to-face interactions. Notably, our results are in line with [47], where the correspondence was examined between data collected by wearable sensors and a video of the same interactions, yielding a sensitivity of 50% (about half of the interactions annotated on the video were present in the wearable sensor data).

Although this limited correspondence between the two methods of measuring interactions could be seen as a negative result, it is striking that, when considering the global social structure extracted from the data, the results of the two methods are in fact strikingly similar. First, at a statistical level, the distributions of events durations are broad in both cases, with most events having a short duration, and a continuously decreasing distribution with no cutoff except the one imposed by the procedure. The distributions of weights (number of events between two individuals) are also very similar. Most importantly, the networks aggregated over the whole observation period turn out to be extremely similar as measured by several metrics: the weights of a link joining two individuals are highly correlated in the two networks, the top ranked links and the strong structures are preserved. Overall, the picture of the social network provided by the two measurement systems are thus extremely similar, despite the discrepancies observed at the very detailed level. We note that this result is at odds with the analysis of [41], in which an interaction network and a proximity network, built from the same set of direct observations of baboons, were shown to differ. However, our infrastructure detects very close proximity, which would be allowed only between animals sharing a certain level of trust, while the proximity criterion used by [41] was of 5 or 10 meters, thus not corresponding to a “contact” between individuals.

Moreover, we have built aggregated networks at shorter timescales and investigated how quickly a reliable network could be obtained in each case. The observation network fluctuates strongly from day to day. It yields a very similar picture with respect to the whole observation period after about 10 days of observation. This is in agreement with recent results showing that limited amounts of observational data was enough to obtain a clear picture of the social structure of the group [22]. Comparatively, the structure of the aggregated network obtained with the sensors is very stable already at short timescales, and we obtain a reliable network structure very similar to the one aggregated over one month even with only one day of data. This implies that the sensor data would potentially allow to pinpoint a change in the structure of the social network of the group on much shorter timescales than with observational data. Moreover, we have actually deployed the sensors infrastructure on much longer timescales with no interruption (currently up to 6 months), while continuous observations cannot be carried out realistically on such timescales.

### Conclusion

Collecting and analyzing data coming from digital devices to evaluate social patterns has become quite common in human social studies. Recently, new infrastructures and protocols based on sensors have also become more easily available for the study of animal groups as an alternative to traditional data collections methods such as behavioral observations. In particular, we have shown the potential of the sensors infrastructure used here, which detects close proximity between individuals, to enable an automatic, less costly, long-term and reliable data collection that yields a picture of the social interactions very similar to the one obtained from direct observations, despite not registering all observed interactions and not distinguishing between different types of behaviors. Such techniques could thus facilitate animal social network analysis and most importantly make accessible both short and long timescales for the investigation of the dynamical evolution of animal social networks [54].

## Acknowledgments

The authors thank the staff at the Rousset-sur-Arc Primate Center (CNRS-UPS846, France) for making the study possible and Julie Gullstrand and Joël Fagot for helping in data collection.

## Data availability

The data that support the findings of this study are openly available in the Open Science Foundation repository at DOI:10.17605/OSF.IO/UFS3Y.

## Authors Contribution

AB and NC designed the study. AB, NC, DP and VG realised the adaptation of the sensors to baboons. VG and JG realized the data collection. VG performed the data analysis and wrote the first draft of the manuscript. AB, NC, VG wrote the final version of the manuscript. All authors gave final approval for publication and agree to be held accountable for the work performed therein.

